# Transfer learning identifies sequence determinants of regulatory element accessibility

**DOI:** 10.1101/2022.08.05.502903

**Authors:** Marco Salvatore, Marc Horlacher, Annalisa Marsico, Ole Winther, Robin Andersson

## Abstract

Dysfunction of regulatory elements through genetic variants is a central mechanism in the pathogenesis of disease. To better understand disease etiology, there is consequently a need to understand how DNA encodes regulatory activity. Deep learning methods show great promise for modeling of biomolecular data from DNA sequence but are limited to large input data for training. Here, we develop ChromTransfer, a transfer learning method that uses a pre-trained, cell-type agnostic model of open chromatin regions as a basis for fine-tuning on regulatory sequences. We demonstrate superior performances with ChromTransfer for learning cell-type specific chromatin accessibility from sequence compared to models not informed by a pre-trained model. Importantly, ChromTransfer enables fine-tuning on small input data with minimal decrease in accuracy. We show that ChromTransfer uses sequence features matching binding site sequences of key transcription factors for prediction. Together, these results demonstrate ChromTransfer as a promising tool for learning the regulatory code.

## Introduction

The human genome encodes hundreds of thousands of transcriptional regulatory elements, including enhancers, promoters, and silencers, that control how genes are expressed in any given cell in the human body (Andersson and Sandelin 2020; Field and Adelman 2020). Regulatory elements are short stretches of DNA that act as regulators of transcription via their ability to interact with key proteins, transcription factors (TFs), that can modulate the expression of genes (Spitz and Furlong 2012; Vaquerizas et al. 2009). Regulatory dysfunction may be caused by disruptions of the regulatory code, for instance through point mutations or structural variants affecting the chromatin accessibility of regulatory element DNA or binding of TFs (Bradner et al. 2017; Miguel-Escalada et al. 2015). Consequently, dysfunction of regulatory elements has emerged as a central mechanism in the pathogenesis of diseases (Zaugg et al. 2022). As a foundation for understanding cellular and disease programs, we therefore need to understand the regulatory code of the human genome. In essence, deciphering genetic variants-to-phenotype associations requires an understanding of how DNA codes for regulatory activities (Andersson and Sandelin 2020; Lappalainen and MacArthur 2021).

The major challenge in understanding the regulatory code is its complexity. Only considering sequences matching known TF binding sequences, regulatory elements involve millions of possible sequences that can encode regulatory function, which can be interpreted differently across cell types. Therefore, experimentally testing every sequence or regulatory element in every cell type is not feasible. Instead, we will need to learn the underlying mechanisms and logic of regulatory elements by building computational models that can be applied to predict regulatory element activity. With an increasing amount of large-scale molecular data for regulatory activities readily available (Andersson et al. 2014; FANTOM Consortium and the RIKEN PMI and CLST (DGT) et al. 2014; Kundaje et al. 2015; Arner et al. 2015; Stunnenberg et al. 2016; Moore et al. 2020; Meuleman et al. 2020), we are now in a position to approach this challenge.

Deep learning approaches show great promise for such a task, due to their ability to detect complex patterns within unstructured data (Ching et al. 2018; Eraslan et al. 2019), as demonstrated by the ability of convolutional neural networks to learn novel features from DNA sequences (Avsec et al. 2021b; de Almeida et al. 2022; Janssens et al. 2022). The major obstacle for their use to derive the regulatory code is the requirement of large input data sets for training the models. Training on small input data may lead to overfitting and, as a consequence, non-generalizable interpretations. Learning efficiency and prediction accuracy can be improved through multi-task learning, in which multiple types of molecular signatures or the same type of measurement across multiple cell types or species (tasks) are modeled jointly through exploitation of commonalities and differences in the data for the different tasks (Eraslan et al. 2019). Such an approach has been successful in genomics, including modeling of chromatin accessibility, histone post-translational modifications, TF binding, and expression from DNA sequence alone (Avsec et al. 2021a; Kelley 2020; Kelley et al. 2018, 2016; Nair et al. 2019; Zhou et al. 2018). However, multi-task learning may lead to optimization imbalances (Chen et al. 2018), causing certain tasks to have a larger influence or even dominate the network weights, which may result in worse accuracy for weaker tasks or inefficacy to separate similar tasks (Avsec et al. 2021a).

Transfer learning (Yosinski et al. 2014) has the potential to avoid the possible problems of optimization imbalances in multi-task learning or overfitting due to small data sets in single-task learning. During transfer learning, a model is first trained on a problem with sufficiently large input data. The knowledge gained during the first stage is then used on a related or more specific problem for which input data may be smaller, using the features and network weights learnt in the first model as a basis for training a new model for the specific problem through fine-tuning. Transfer learning has been highly successful in biological image classification (Esteva et al. 2017; Zeng et al. 2015) and also shows great promise for training models from DNA sequences to predict 3D genome folding (Schwessinger et al. 2020), chromatin accessibility (Nair et al. 2019; Kim et al. 2021; Kelley et al. 2016; Lai et al. 2022), and TF binding (Zheng et al. 2021; Novakovsky et al. 2021).

As a step towards establishing transfer learning for modeling the regulatory code, we here develop ChromTransfer, a transfer learning scheme for single-task modeling of the DNA sequence determinants of regulatory element activities (Figure 1A). ChromTransfer uses a pre-trained, cell-type agnostic model, that we derive from a large compendium of permissively defined regulatory elements from open chromatin regions (Figure 1B) across human cell types, tissues, and cellular stages, to fine-tune models for specific tasks (Figure 1C). We demonstrate improvements in performances with ChromTransfer for predicting cell-type specific chromatin accessibility for all cell types considered compared to baseline models derived from direct modeling of individual cell types. We find that transfer learning minimizes overfitting, allowing fine-tuning of models with high predictive performances using only a small fraction of input data. Through feature importance analysis, we identify how ChromTransfer uses sequence elements to predict chromatin accessibility differently across cell types and match these elements to key TF binding site sequences. Our results demonstrate ChromTransfer as a promising tool for deciphering how DNA codes for regulatory activities from small input data.

**Figure 1:**
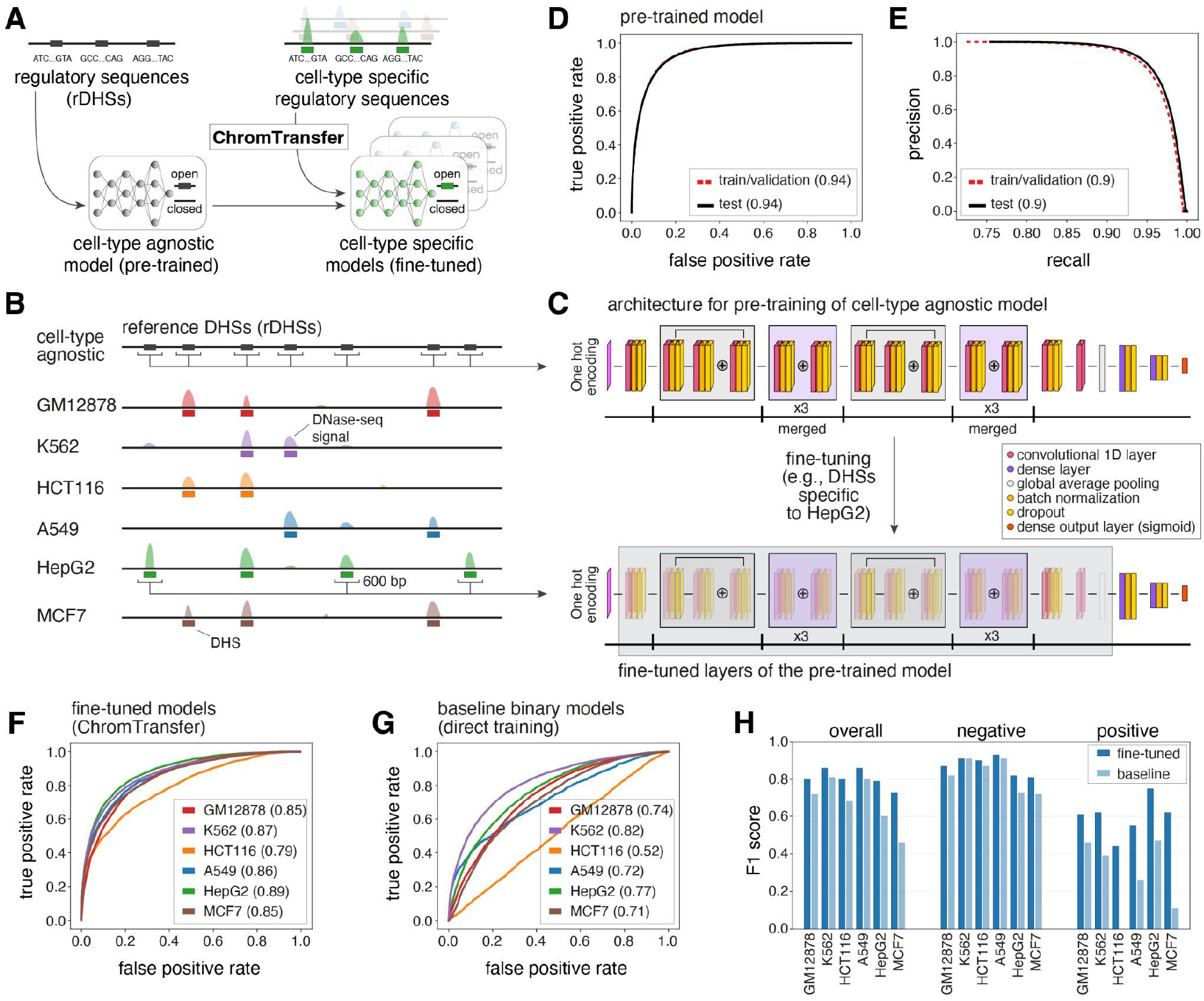
Transfer learning of the sequence determinants of regulatory elements using ChromTransfer. **A**: ChromTransfer is a transfer learning scheme for single-task modeling of the DNA sequence determinants of regulatory element activities. ChromTransfer uses a pre-trained, cell-type agnostic model, derived from a large compendium of open chromatin regions to fine-tune models for predicting cell-type specific activities. **B**: Illustration of a genomic locus with DNase-seq signal across six cell lines along with called DHSs and the cell-type agnostic rDHS compendium. The strategy for selection of positives, 600 bp sequences centered on all rDHSs (for pre-training) or cell-type specific DHSs (for fine-tuning) are shown. **C**: Model architecture (upper panel) and strategy for fine-tuning (lower panel). For network details, see Methods. **D**: ROCs for training/validation and the test set of the pre-trained model for rDHS classification. AUROCs are provided in parentheses. **E**: Precision recall curves (PRCs) for training/validation and the test set for the pre-trained model for rDHS classification. AUPRCs are provided in parentheses. **F-G**: Test set ROCs of the six fine-tuned models (F, ChromTransfer) and the six binary class baseline models (G, direct training scheme) for classification of cell-type specific chromatin accessibility. AUROCs for each cell line model are provided in parentheses. **H**: Overall and per-class (positive: open chromatin, negative: closed chromatin) test set F1 scores for the fine-tuned and binary class baseline models of the six considered cell lines. F1 scores are also given in Supplementary Table 1.

## Results

### Transfer learning improves regulatory element prediction accuracy compared to direct learning

As a basis for learning sequence features associated with chromatin accessibility, we considered the ENCODE compendium of 2.2 million representative DNase I hyper-sensitive sites (rDHSs, cell-type agnostic open chromatin regions, https://screen.encodeproject.org/) (Moore et al. 2020) (Figure 1B). The ENCODE rDHSs were assembled using consensus calling from 93 million DHSs called across a wide range of human cell lines, cell types, cellular states, and tissues, and are therefore likely capturing the great majority of possible sequences associated with human open chromatin.

**Table 1:** ChromTransfer allows fine-tuning on small input data. Mean overall and per-class (positive: open chromatin, negative: closed chromatin) F1 scores on test set data for HepG2 models fine-tuned after bootstrapping training data (1% to 75% of original training data, 10 bootstraps). The overall F1 score of the original HepG2 fine-tuned model (100%) is included for reference. Bootstrap sample sizes refers to the total number of training examples, including both positive and negative examples. Standard deviations (sd) across bootstrap estimates are included.

| Sample (%) | Sample (size of training) | mean F1 negative | sd F1 negative | mean F1 positive | sd F1 positive | mean F1 overall | sd F1 overall |
| --- | --- | --- | --- | --- | --- | --- | --- |
| 1 % | 747 | 0.65 | 0.22 | 0.68 | 0.03 | 0.67 | 0.12 |
| 5 % | 3733 | 0.76 | 0.01 | 0.72 | 0.01 | 0.74 | 0.01 |
| 10 % | 7467 | 0.77 | 0.01 | 0.71 | 0.04 | 0.74 | 0.02 |
| 15 % | 11201 | 0.76 | 0.02 | 0.73 | 0.01 | 0.75 | 0.01 |
| 20 % | 14935 | 0.77 | 0.01 | 0.74 | 0.01 | 0.75 | 0.01 |
| 25 % | 18668 | 0.77 | 0.01 | 0.74 | 0.02 | 0.75 | 0.01 |
| 50 % | 37366 | 0.79 | 0.01 | 0.76 | 0.01 | 0.77 | 0.01 |
| 75 % | 56005 | 0.80 | 0.01 | 0.77 | 0.01 | 0.79 | 0.01 |
| 100 % | 74673 | 0.82 | - | 0.75 | - | 0.79 | - |

We implemented a ResNet (He et al. 2015) inspired deep neural network architecture with residual layers to classify chromatin accessibility (open/closed) from 600 bp DNA sequences centered at rDHSs (Figure 1C, upper panel). The network was used for cell-type agnostic modeling of chromatin accessibility versus sampled negative genomic regions (Methods). Training and hyperparameter tuning were carried out using 3-fold cross-validation. rDHSs located on chromosomes 2 and 3 were held out as the test set. The resulting model (herein referred to as pre-trained) was capable of distinguishing between open and closed chromatin with high accuracy (area under receiving operating curve (AUROC) of 0.94 and area under precision-recall curve (AUPRC) of 0.90 for the out of sample test set; per-class test set F1 scores of 0.93 and 0.80 for open and closed chromatin, respectively; Figure 1D,E). This demonstrates that DNA sequence is a major determinant of chromatin accessibility, in agreement with previous work (Kelley et al. 2016; Nair et al. 2019), and that the pre-trained model is able to capture the high sequence complexity in the input data.

To examine how well the pre-trained model could be transferred to more specific prediction tasks with limited training data, we developed a new transfer learning procedure, ChromTransfer, and evaluated its ability to learn the sequence determinants of open chromatin regions unique to specific cell types (Figure 1A-C). We focused on rDHSs with cell-type specific chromatin accessibility across six cell lines (defined as accessible sites unique to one cell line among the six considered cell lines; Figure 1B; GM12878: 31,740, K562: 36,769, HCT116: 20,018, A549: 14,112, HepG2: 31,211, MCF7: 39,461) together reflecting diverse biological cell types, each with its own key TFs (The ENCODE Project Consortium 2012). During transfer learning with ChromTransfer, the representations of the higher-order features in the convolutional blocks of the pre-trained model are re-trained to make them more relevant for the new data alongside training of newly added dense layers. In order to gradually adapt the pre-trained features to the new data, both the convolutional blocks and the dense layers are trained at a reduced learning rate (Figure 1C, lower panel; Methods). In this way, the pre-trained model can be fine-tuned to capture the sequence determinants of chromatin accessibility in individual cell types. A similar approach was taken to fine-tune a general model to capture time-point specific chromatin accessibility during epidermal differentiation (Kim et al. 2021).

ChromTransfer achieved high predictive performances for all cell lines (overall test set F1 scores ranging between 0.73 and 0.86, AUROC ranging between 0.79 and 0.89, and AUPRC ranging between 0.4 and 0.74; Figure 1F; Supplementary Figure 1B; Supplementary Tables 1-2). In comparison, the pre-trained model (without fine-tuning) demonstrated only a weak ability to predict cell-type specific chromatin accessibility (overall test set F1 scores ranging between 0.24 and 0.49; Supplementary Table 1), indicating that fine-tuning of the pre-trained model adapts the network weights to capture cell-type specific sequence elements. The largest improvement was observed for K562, having an increase in overall F1 score from 0.24 for the pre-trained model (per class test set F1 score of 0.22 and 0.33 for closed and open chromatin, respectively) to 0.86 for the fine-tuned model (per class test set F1 score of 0.91 and 0.62 for closed and open chromatin, respectively).

We next examined if transfer learning using ChromTransfer added any performance increase compared to a direct training approach. As baseline models, we trained the same ResNet-like network (Figure 1C, upper panel) *ab initio* using the same DNA sequences from cell-type specific open chromatin regions for each of the six cell lines either separately as a binary classification task (one model per cell type) or together with a multi-class classification output layer (one model for all six cell types). Hence, any performance differences observed for the binary class baseline models versus ChromTransfer models will reflect the absence of cell-type agnostic pre-training of the convolutional layers on the rDHSs. Indeed, the fine-tuned models consistently outperformed the direct binary class training scheme (mean increase in overall test set F1 score of 0.13, ranging between 0.05 for K562 to 0.27 for MCF7; Figure 1G,H; Supplementary Figure 1B,C; Supplementary Table 1). The largest performance increase was observed for the positive class (open chromatin), with HCT116 and MCF7 binary class baseline models having very weak positive predictive performances (test set positive class F1 score of 0 and 0.11, respectively). Similarly, the fine-tuned ChromTransfer models demonstrated consistent improvements in predictive performances over the multi-class baseline model (mean increase in overall test set AUROC of 0.15; Supplementary Table 3).

We conclude that ChromTransfer’s pre-training of a cell-type agnostic model on the sequence determinants of chromatin accessibility followed by fine-tuning on individual cell-types consistently improves classification accuracy.

### Transfer learning allows for fine-tuning on small training data without overfitting

The weak class performances for some cell line models with the direct training scheme (baseline models; Figure 1H; Supplementary Table 1) indicates that training of the network architecture on these data sets is not capable of generalizing to the test data. Indeed, examination of the learning curves (Figure 2A) showed clear signs of overfitting to the training data, with early stopping only after a few epochs and limited convergence between validation and training losses. Further examination revealed that the direct training scheme for the binary class baseline models could not properly calibrate class probabilities (lack of external calibration; Figure 2C; Supplementary Figure 2). In contrast, the ChromTransfer-derived fine-tuned models showed no signs of overfitting (Figure 2B,D).

**Figure 2:**
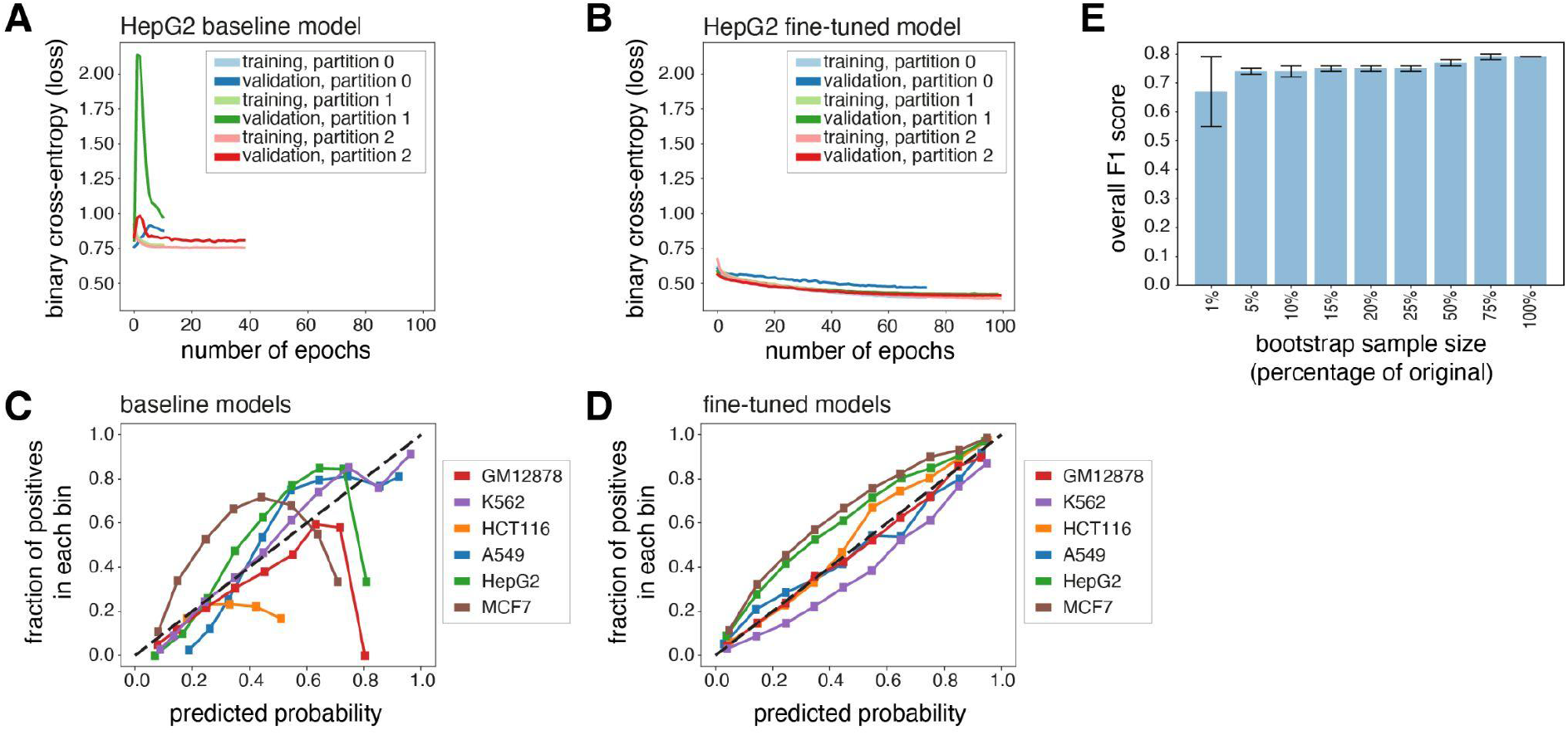
Transfer learning minimizes overfitting and enables training on small data sets. **A-B**: Learning curves (binary cross-entropy, loss) for training and validation data for the HepG2 binary class baseline model (direct training scheme; A) and fine-tuned model (ChromTransfer; B) for 3-fold cross-validation (partitions 0, 1, 2). **C-D**: Calibration curves displaying the observed fraction of positives (vertical axes) across 10 bins of predicted probabilities of positive class (horizontal axes) for unseen test set data (external calibration) for each of the six binary class baseline models (direct training scheme; C) and the six fine-tuned models (D; ChromTransfer). The number of samples in each predicted probability bin and the bin ranges for each cell line and model are shown in Supplementary Figure 2. **E**: Overall F1 scores on test set data for HepG2 models fine-tuned after bootstrapping training data (1% to 75% of original training data). The overall F1 score of the original HepG2 fine-tuned model is included for reference. Error bars show standard deviations (10 bootstraps). For exact values, see Table 1.

The stable performances of ChromTransfer’s fine-tuned models without indications of overfitting prompted us to investigate how small training datasets could be used without a major decline in predictive performance. To this end, we performed a bootstrap analysis in which we subsampled the training data for HepG2-specific chromatin accessibility to different fractions (1% to 75%, 10 bootstraps per target) of the original training data and re-ran the fine-tuning of the pre-trained model on the resulting data (Table 1). Remarkably, we observed only a marginal decrease in predictive performance (decrease in mean overall test set F1 score of 0.05) on the original test data when using as low as 5% of the training data (3,733 input sequences, among which 1,283 were positive training examples; Figure 2E). With only 1% of the training data (747 sequences, 257 positives), we observed a slightly larger reduction (mean overall test set F1 score of 0.67) and more variation (test set overall F1 score standard deviation of 0.12) in performances. Still, the models fine-tuned on 1% of the training data outperformed both the pre-trained model (overall test set F1 score of 0.49) and the binary class baseline model derived from the direct training scheme (overall test set F1 score of 0.60).

Taken together, we conclude that ChromTransfer allows for accurate sequenced-based modeling of chromatin accessibility using small input data sets while being robust towards overfitting. This suggests that ChromTransfer-derived models capture the regulatory code for chromatin accessibility.

### Feature importance analysis reveals the importance of TF binding site sequences for the fine-tuned models

To investigate the underlying sequence patterns used by ChromTransfer when making predictions, we performed feature importance analysis using gradient × input (Eraslan et al. 2019; Shrikumar et al. 2019). We specifically focused on how the importance of individual base pairs had changed during fine-tuning. To this end, we focused on 27,940 and 35,179 positive predictions by the HepG2 and K562 fine-tuned models, respectively. HepG2 cells are derived from a hepatocellular carcinoma, while K562 cells are of erythroleukemia type. These two cell lines are therefore expected to have highly different regulatory activities and active TFs.

Feature importance analysis revealed both increased and decreased importance for individual base pairs in the fine-tuned HepG2, compared to the pre-trained model. This is exemplified by increased importance of sequences at putative binding sites for HNF4A and HNF4G (Figure 3A), hepatocyte nuclear factors, and CEBPA and CEBPD (Figure 3B), CCAAT/enhancer-binding proteins (CEBPs), all of critical importance for hepatocyte function and differentiation (Akai et al. 2014; Hayhurst et al. 2001). In contrast, we observed a decreased importance for sequences matching binding sites of non-hepatocyte TFs, for instance OLIG2, NEUROD2 and TAL1-TCF3 (Figure 3C). OLIG2 and NEUROD2 are important for the central nervous system and neurodevelopment (Takebayashi et al. 2000; Olson et al. 2001) while TAL1-TCF3 is required for early hematopoiesis (Hoang et al. 2016), and neither are likely to be important for hepatocytes. This indicates that transfer learning can refocus on relevant sequence elements important for the task at hand.

**Figure 3:**
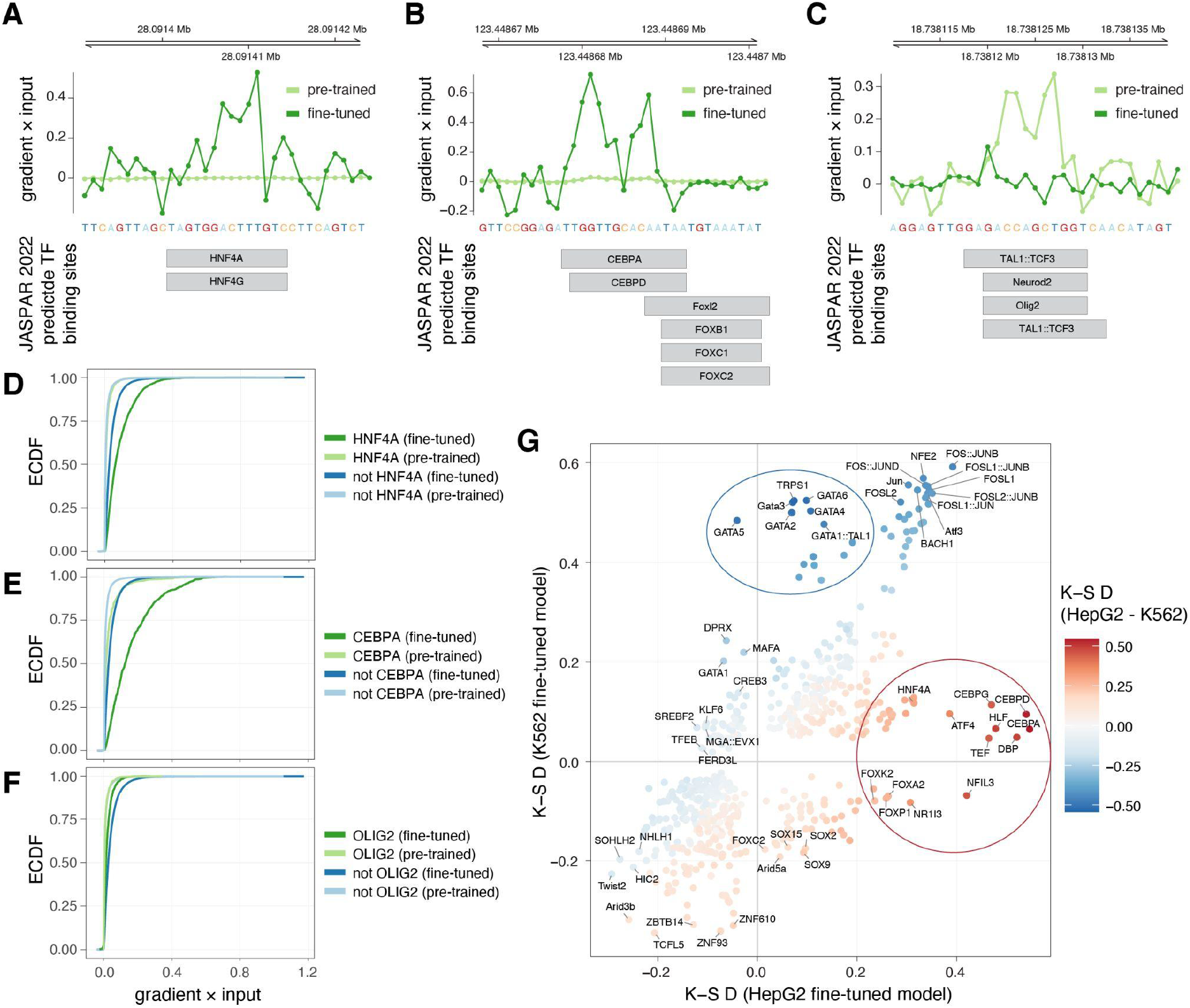
Feature importance analysis reveals TFs important for cell-type specific chromatin accessibility of regulatory elements. **A-C**: Feature importance scores (gradient × input, upper panels) at example loci overlapping JASPAR 2022 predicted TF binding sites (lower panels), highlighting increased importance of base pairs at putative binding sites for HNF4A/G (A) and CEBPA/D (B), and decreased importance of base pairs at those for NEUROD2, OLIG2, and TAL1-TCF3 heterodimer. (C). **D-F**: Empirical cumulative distribution functions (ECDF, vertical axes) of feature importance scores (gradient × input, horizontal axes) associated with predicted binding sites of HNF4A (D), CEBPA (E), and OLIG2 (F) in fine-tuned and pre-trained models. ECDFs for that of predicted binding sites for all other TFs not overlapping target TFs (HNF4A, CEBPA, or OLIG2) are shown for comparison. **G**: Kolmogorov-Smirnov (K-S) test statistics (D statistics) for feature importance scores (gradient × input) associated with predicted binding sites of each considered TF in the HepG2 (horizontal axis) and K562 (vertical axis) fine-tuned models. TFs are colored according to a KS D statistic calculated from the differences between the importance of TFs for the two models based on their association with gradient × input scores. Only TFs with Benjamini-Hochberg adjusted FDR < 0.001 are shown. TFs of biased importance for HepG2 and K562 models are highlighted with red and blue ellipses, respectively.

To systematically evaluate sequence elements important for the two fine-tuned models, we overlaid the feature importance scores with predicted binding sites from the JASPAR 2022 motif database (Castro-Mondragon et al. 2022) and associated each predicted TF binding site with the max corresponding score (Methods). Examination of the distributions of the feature importance scores for individual TFs versus all TFs considered confirmed the individual observations above, with overall high importance for HNF4A and CEBPA in the HepG2 model (Figure 3D,E) and low importance for OLIG2 (Figure 3F), while these rank differences were not observed for the pre-trained model. In-silico mutagenesis (ISM) delta scores were in large agreement with the feature importance scores derived from gradient × input (Supplementary Figure 3).

Based on these observations, we calculated Kolmogorov-Smirnov (K-S) rank statistics (D statistics) to examine differences between the importance of TFs for the models based on their association with gradient × input scores. This analysis revealed major differences between the two fine-tuned models. CEBPs were ranked among the most important TFs for HepG2 cells, while GATA factors, critical for the development and maintenance of the hematopoietic system (Gao et al. 2015), were ranked among the most important TFs for K562 cells (Supplementary Figure 4). Furthermore, both CEBP and GATA factors displayed increased importance in the respective fine-tuned models compared to the pre-trained model (Supplementary Figure 5).

Direct comparison between the two fine-tuned models (Figure 3G) highlighted GATA factors (K562), CEBPs (HepG2), and HNF4A (HepG2), alongside Forkhead box proteins, DPB, HLF, NFIL3, and TEF (HepG2) as the most discerning TFs for the two cell lines. Although PAR bZIP (proline-and acid-rich basic region leucine zipper) TFs NFIL3, DBP, TEF, and HLF all recognize similar binding site sequences, similar to the ambiguity between HNF4A and HNF4G and that of CEBPA and CEBPD (Castro-Mondragon et al. 2022), making it hard to predict actual TF binding, they are all of relevance for hepatocyte function (Mueller et al. 1990; Cowell and Hurst 1996). FOS-JUN heterodimer binding site sequences were, on the other hand, found important for both cell lines (Figure 3G; Supplementary Figure 4), and had an increased importance compared to the pre-trained model (Supplementary Figure 5).

These results demonstrate that transfer learning from a pre-trained model derived from a large compendium of DHSs based on cell-type specific regulatory elements with ChromTransfer does not only yield improved prediction accuracy, but also reveals the underlying sequence elements of relevance for the regulatory elements, indicating that ChromTransfer has a large potential to further our understanding of the regulatory code.

## Discussion

The major challenge in understanding the regulatory code is its complexity. Only considering sequences matching known TF binding sequences, regulatory elements involve millions of possible sequences that can encode regulatory function, which can be interpreted differently across cell types. Therefore, experimentally testing every sequence or regulatory element in every cell type is not feasible. Instead, we will need to learn the underlying mechanisms and logic of regulatory elements by building computational models that can be applied to predict regulatory element activity.

Understanding the regulatory code will be transformative for the field, ultimately allowing direct interpretation of disease-associated genetic variants, fine-mapping of risk alleles, and a direct interpretation of cell types involved in disease etiology. However, computational modeling of the regulatory code has been hampered by the requirement of large data sets for training, especially for deep learning (Ching et al. 2018; Eraslan et al. 2019), and failure to meet this requirement may lead to non-generalizable models. We here establish a transfer learning scheme, ChromTransfer, that exploits available large-scale data sets for training of a general sequence model of regulatory elements that can be fine-tuned on a specific problem for which only a small amount of data is available or can be generated. As a proof-of-concept, we demonstrate that this approach is insensitive to overfitting, even at minuscule data sizes, allowing accurate modeling of the sequence determinants of cell-type specific chromatin accessibility. In contrast, using the same network architecture trained *ab initio* on the same data failed to produce generalizable results, indicating that transfer learning is required for such a modeling task, at least with the current network architecture. We note that the amount of data required for fine-tuning will depend on the complexity of the prediction task at hand. A higher sequence complexity underlying cell-type specific chromatin accessibility, enabling binding of different proteins alone or in combinations, likely explains why transfer learning for this task still requires larger training data than learning the binding of individual TFs from sequence (Novakovsky et al. 2021).

For ease of validation, we here focused on well-studied cell lines with known master regulatory TFs. Feature importance analysis using gradient × input revealed binding site sequences for these key TFs to be most important for predicting cell-type specific chromatin accessibility, which were further supported by in-silico mutagenesis. Although our analysis shows promise, establishing the regulatory code will require broad analysis across multiple cell types and more in-depth modeling of different regulatory activities, e.g., enhancer versus promoter function (Andersson and Sandelin 2020), as well as context and stage-specific activities. We expect that such efforts should be feasible with ChromTransfer. ChromTransfer models may, for instance, enable investigations of the mechanisms underlying dynamic activities during development and in response to cellular stimuli. Such questions are frequently limited to few data points, tasks which are suitable for transfer learning. We further acknowledge that further work on model interpretation is needed to arrive at a sequence code for regulatory activity. Recent developments to this end (Shrikumar et al. 2019; Avsec et al. 2021b; de Almeida et al. 2022; Taskiran et al. 2022) show great promise, and we expect that integration of such analyses with the transfer learning scheme of ChromTransfer will be important for future efforts to understand the regulatory code.

## Methods

### Data used for modeling of regulatory sequences

We considered the ENCODE compendium of 2.2 million rDHSs (Moore et al. 2020) as positives for training a cell-type agnostic neural network (pre-trained model) the sequence determinants of chromatin accessibility. The rDHSs were originally derived from 93 million DHSs called by ENCODE (Moore et al. 2020) and the Roadmap Epigenomics (Kundaje et al. 2015) projects from hundreds of human biosamples, including cell lines, cell types, cellular states, and tissues, and was originally used as an entry point for in the ENCODE Registry of candidate cis-regulatory elements (cCREs, described at https://screen.encodeproject.org/). For each rDHS, we extracted the plus strand 600 bp sequence (GRCh38) centered on the rDHS midpoint. 600 bp was used to make sure that sequences influencing regulatory activity and chromatin accessibility contained within a central open chromatin site (150-300bp) as well as within flanking nucleosomal DNA (150-200bp) were captured (FANTOM Consortium and the RIKEN PMI and CLST (DGT) et al. 2014; Meuleman et al. 2020). To ensure that the pre-trained model was not biased towards any of the specific cell lines considered beforehand, all rDHSs with called accessibility in any of the considered six cell lines (described below) were removed before training of the pre-trained model. This is not necessary for the modeling purpose per se, but was done to test ChromTransfer’s ability to fine-tune the pre-trained model to new, unseen data. Negatives were derived from tiling the genome (GRCh38) in 600 bp non-overlapping windows using BedTools (Quinlan and Hall 2010), followed by removal of any region that overlapped gaps in the GRCh38 genome assembly or manually curated ENCODE blacklist regions (ENCFF356LFX) (Amemiya et al. 2019; The ENCODE Project Consortium 2012), or those within 300 bp of rDHSs.

For modeling of cell-type specific chromatin accessibility (fine-tuning), we considered human cell lines A549, HCT116, HepG2, GM12878, K562, and MCF7. The chromatin accessibility of each rDHSs in each of these cell lines were quantified as described elsewhere (Moore et al. 2020). In summary, ENCODE BigWig signals were aggregated in each rDHS for each replicate of the cell line, followed by a global Z-score transformation of the log_10_-transformed signal aggregates. Z-scores were binarized into closed/open using a threshold of 1.64. Finally, rDHSs were considered open if they were called open in any replicate of the cell line. We defined positives for each cell line as rDHSs that were only accessible in that cell line among the six cell lines considered (GM12878: 31,740, K562: 36,769, HCT116: 20,018, A549: 14,112, HepG2: 31,211, MCF7: 39,461), while negatives (GM12878: 81,805, K562: 103,995, HCT116: 62,389, A549: 78,725, HepG2: 54,995, MCF7: 91,122) were sampled from the positives of the other cell lines and the rDHSs used for pre-training (positive:negative ratio ranging between 1:2.5 and 1:3.5).

### Neural network architecture, training and hyperparameter tuning of the pre-trained model

We implemented a ResNet (He et al. 2015) inspired neural network, visualized in Figure 1C (upper panel). The neural network model uses as input one-hot-encoded DNA sequences (A = [1,0,0,0], C = [0,1,0,0], G = [0,0,1,0], T = [0,0,0,1]) of 600 bp to predict closed (negative) or open (positive) chromatin as output. The neural network consists of a 1-dimensional convolutional layer with 64 hidden units and a kernel size of 25, followed by a residual block with 32 hidden units and a kernel size of 20, 3 merged blocks without residual connections with 32 hidden units and a kernel size of 15, another residual block with 64 hidden units and a kernel size of 10, another 3 merged blocks without residual connections with 64 hidden units and a kernel size of 5, two 1-dimensional convolutional layers with 64 hidden units each and a kernel size of 10 and 5, global average pooling and two dense layers with 512 and 128 nodes. Batch normalization and dropout (0.1) were applied after each layer. The activation function ReLU (Agarap 2019) was used in all layers except the last, in which a sigmoid activation function was used to predict the final class (negative or positive).

rDHSs located on chromosomes 2 and 3 were only used as the test set and rDHSs from the remaining chromosomes were used for training and hyperparameter tuning with 3-fold cross-validation. Hyperparameters were adjusted to yield the best performance on the validation set. The neural network model was implemented and trained in Keras (version 2.3.1, https://github.com/fchollet/keras) with the TensorFlow backend (version 1.14) (Abadi et al. 2016) using the Adam optimiser (Kingma and Ba 2017) with a learning rate of 0.001, batch size of 256, and early stopping with a patience of 15 epochs. Both pre-trained and fine-tuned models (see below) were trained on a Linux SMP Debian 4.19.208-1 x86_64 machine using NVIDIA Quadro RTX 6000 cards with 24 GB of VRAM.

### Training and hyperparameter tuning of the fine-tuned models

For fine-tuning of the pre-trained model, the trained convolutional blocks of the pre-trained model were transferred to a new model and the last two dense layers of the network were adjusted to 1024 and 32 nodes, respectively, and added anew. Batch normalization and dropout (0.1) were applied after each layer. As in the pre-training phase, rDHSs from chromosomes 2 and 3 were only used as a test set for the fine-tuned models. The regions of the remaining chromosomes were used for training and tuning of the hyperparameters with 3-fold cross-validation. The hyperparameters were tuned to give the best performance in the validation set. In order to gradually adapt the pre-trained features to the new data, both the convolutional blocks and the dense layers are trained with a considerably lower learning rate (0.000005). Training was performed with a batch size of 128 and early stopping with a patience of 10 epochs. Time per epoch was 142s (2ms/sample) for a total ∼82K samples (A549) and 60 epochs (142 minutes, average of the 3 cross validation models).

### Training and hyperparameter tuning of the binary and multi-class baseline models

For training and fine-tuning of the binary class baseline models we used the same architecture as when training the pre-trained model (see above), but adjusted the last two dense layers before the output layer of the network to 1024 and 32 nodes, respectively. As in the pre-training and fine-tuning phases, rDHSs from chromosomes 2 and 3 were only used as a test set for the baseline models. The regions of the remaining chromosomes were used for training and tuning of the hyperparameters with 3-fold cross-validation. The hyperparameters were tuned to give the best performance in the validation set. Training was performed with a learning rate of 0.001, batch size of 128, and early stopping with a patience of 10 epochs. Time per epoch was 142s (2ms/sample) for a total ∼82K samples (A549) and 100 epochs (236 minutes, average of the 3 cross validation models).

Lastly, we trained a multi-class model as an additional baseline model, using the same architecture of the fine-tuned and binary class baseline models, except for the output layer, which was changed to a fully connected layer with 7 units (one for each cell line and one for the universal negative class). The model was trained for 50 epochs on the concatenation of cell-line specific fine-tuning datasets. Due to differences in class frequencies when compared to the binary classification models, performance of the multi-class model was evaluated via the class balance agnostic ROC curve metric (AUROC).

### Bootstrap analysis and evaluation of overfitting

To examine the impact of training data size on model fine-tuning, we performed a bootstrap analysis using HepG2 training data. The original training data of 74,673 input sequences were subsampled to target sizes of 1%, 5%, 10%, 15%, 20%, 25%, 50%, and 75% in 10 bootstraps each, followed by fine-tuning of the pre-trained model (as above). For each bootstrap, the overall and per-class F1 scores for the test set (chromosome 2 and 3) were calculated and the mean and standard deviations of F1 scores were reported for each target size.

Overfitting of the models was evaluated by inspection of the cross-validation and training accuracies using learning curves of the validation and training losses (binary cross-entropy). We further evaluated the disagreement between observed and predictive probabilities by inspection of calibration curves on the test data (external calibration).

### Feature importance analysis

To investigate the sequence elements underlying the predictions of the pre-trained model and the K562 and HepG2 fine-tuned models, we calculated feature importance scores as the dot product between the input DNA sequence gradients (with respect to the output neuron) and the one-hot encoding of the sequence (gradient × input). For comparison with the pre-trained model, feature importance scores were derived from the 27,940 and 35,179 positive predictions (from training and test data) of the HepG2 and K562 models respectively. For comparison between the K562 and HepG2 fine-tuned models, we considered the union (62,689) of positive predictions.

Since gradient × input scores may have problems to correctly estimate the importance of sequence elements for making predictions in case of multiple occurrences in the same input sequence (Eraslan et al. 2019; Shrikumar et al. 2019), we evaluated their agreement with delta scores derived from in-silico mutagenesis (ISM). For computational reasons, we limited the ISM calculations to true positives (from training and test data) of the HepG2 and K562 fine-tuned models. For each input sequence and base pair, we calculated the max difference in output probability (ISM delta score) after mutating the original nucleotide to any of the other three nucleotides. Nucleotides within the original input sequences important for positive predictions of a model will yield a negative ISM delta score (decrease in positive prediction score) in contrast to gradient × input scores of important sequences that will be positive. Validation of gradient × input scores by ISM delta scores were performed for the HepG2 model by correlating the feature importance scores associated with predicted TF binding sites (see below) of CEBPA, HNF4A, and FOS-JUNB heterodimer.

### Model interpretation using predicted TF binding sites

To systematically evaluate sequence elements important for the two fine-tuned models, we analyzed the gradient × input and ISM delta scores with respect to predicted binding sites from the JASPAR 2022 motif database (derived from motif scanning; P < 1e-5) (Castro-Mondragon et al. 2022) using R (version 4.0.3) (R Core Team 2022). Results were plotted using ggplot2 (version 3.3.5) (Wickham 2016) and Gviz (version 1.34.1) (Hahne and Ivanek 2016). Predicted TF binding sites were imported and overlaid rDHS regions using rtracklayer (version 1.55.4) (Lawrence et al. 2009) and GenomicRanges (version 1.42.0) (Lawrence et al. 2013). Each predicted TF binding site was associated with the maximum gradient × input score (or minimum in-silico mutagenesis delta score) across the contained base pairs. Only TFs with at least 100 predicted binding sites across all considered rDHSs were considered.

The importance of each TF for each model was evaluated through rank-based enrichments of the importance scores of its predicted rDHS-associated binding sites versus the importance scores of non-overlapping predicted binding sites of all other TFs. Evaluation was carried out using both manual inspection of the associated empirical cumulative distribution functions and, more systematically, using the Kolmogorov-Smirnov test. For visualization purposes, we changed the sign of the resulting D statistic if the average ranks of the scores for the predicted binding sites of a TF were smaller than the average ranks of all other non-overlapping binding sites. Only TFs with significant deviation from the null hypothesis of no difference in rank (Benjamini-Hochberg adjusted FDR < 0.001) were plotted.

## Code availability

Code for modeling, feature attribution analysis and model interpretation performed in this study, as well as trained models are publicly available on GitHub: https://github.com/anderssonlab/ChromTransfer/ (v1.1). The repository has been archived at Zenodo (https://doi.org/10.5281/zenodo.7528544).

## Acknowledgements

We thank members of the Andersson lab and Winther lab for rewarding discussions. This work was supported by funding from the Novo Nordisk Foundation [grants NNF18OC0052570, NNF20OC0059796, NNF21SA0072102 to R.A. and NNF20OC0062606 to O.W.]. M.H. was supported by the Helmholtz Association under the joint research school “Munich School for Data Science - MUDS”.

## Author contributions

M.S. and R.A. conceived the project; M.S. and M.H. performed machine learning and feature attribution analysis with support from O.W., A.M., and R.A.; R.A. performed downstream model interpretation analyses; R.A. supervised the project; R.A. wrote the manuscript with input from M.S., M.H., A.M., and O.W.; all authors reviewed the final manuscript.

