## Supplementary Figures 1-5 and Supplementary Tables 1-3 for "Transfer learning identifies sequence determinants of regulatory element accessibility"

* Shared first authors

**Contents**

Supplementary Figures 1-5

Supplementary Tables 1-3

### Supplementary Figures

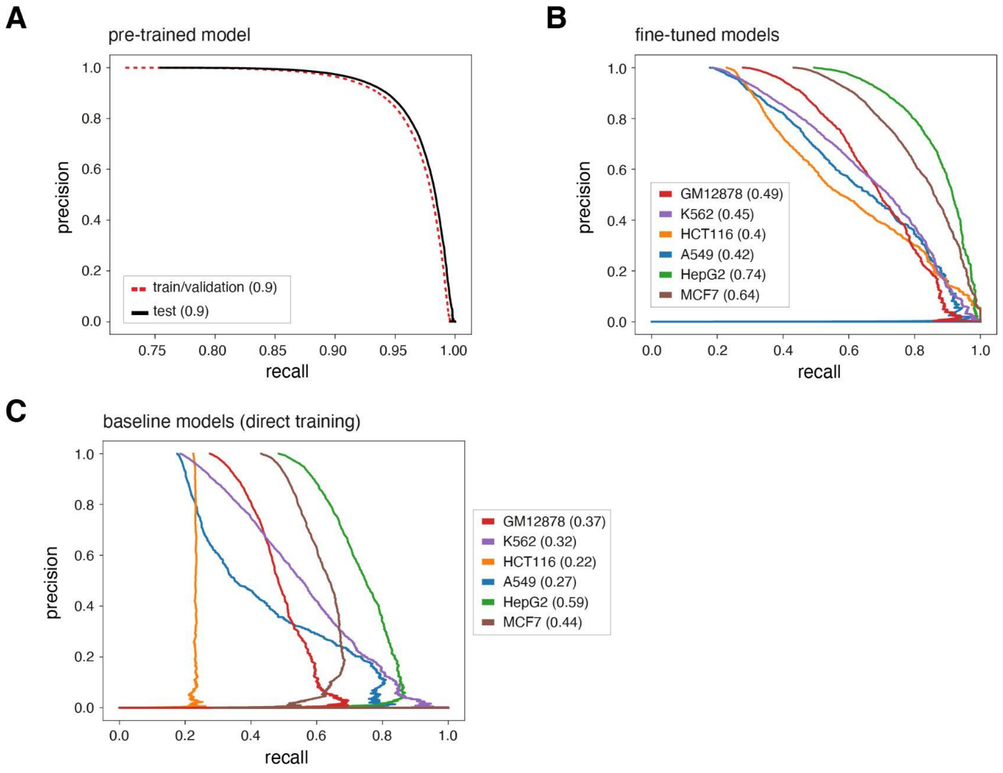


**Supplementary Figure 1: Predictive performances of the models. A**: Precision recall curves (PRCs) for training/validation and the test set for the pre-trained model for rDHS classification. AUPRCs are provided in parentheses. **B**: PRCs for the six ChromTransfer fine-tuned models for classification of cell-type specific chromatin accessibility. AUPRCs for each cell line are provided in parentheses. **C**: PRCs for the six binary class baseline models (direct training scheme) for classification of cell-type specific chromatin accessibility. AUPRCs for each cell line are provided in parentheses.

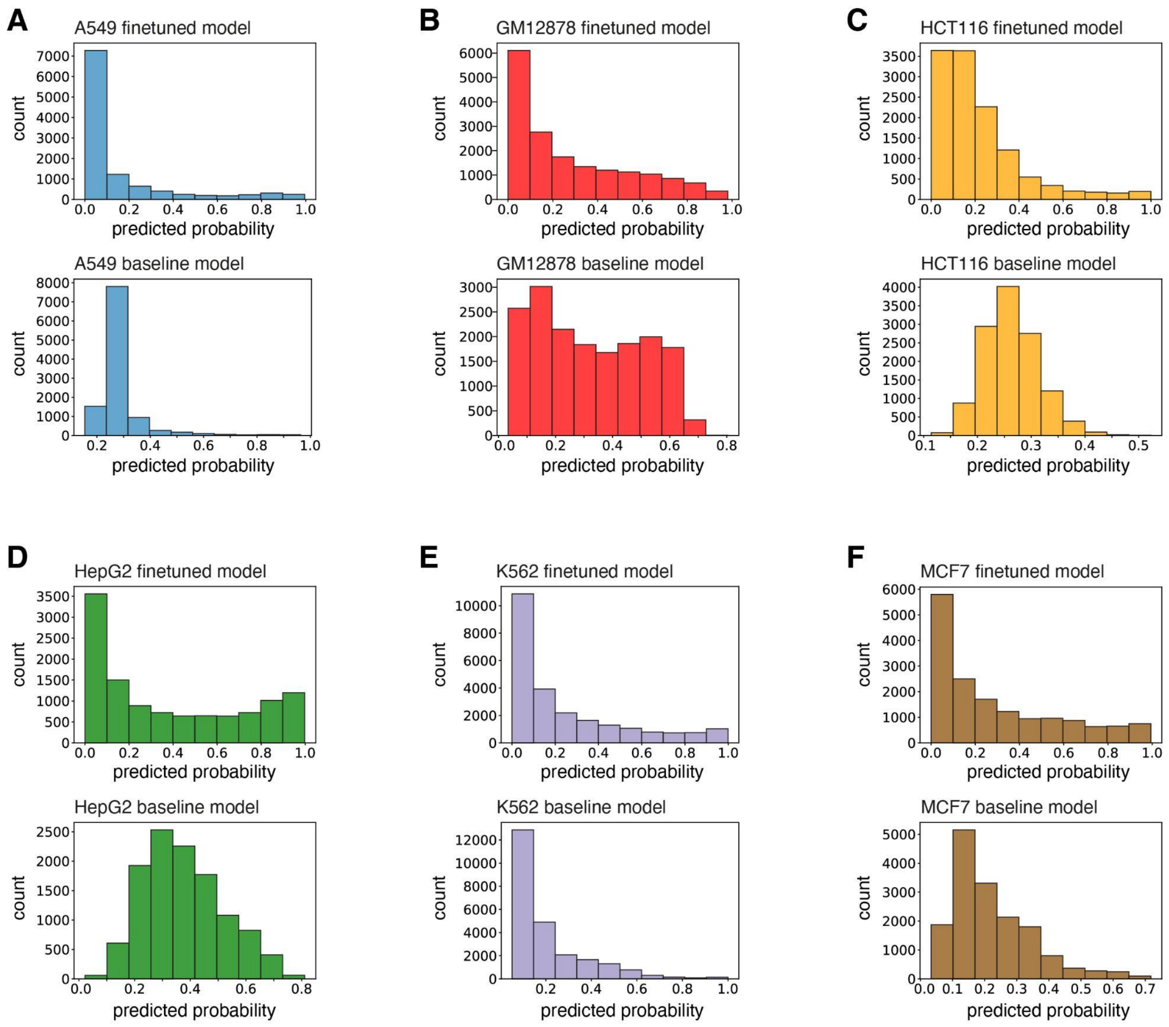


**Supplementary Figure 2: Sample counts per predicted probability bin. A-F**: The number of samples in each predicted probability bin (vertical axes) and the distributions of bins (horizontal axes, 10 bins) for cell lines A549 (A), GM12878 (B), HCT116 (C), HepG2 (D), K562 (E), and MCF7 (F). For each cell line: top panel displays the sample count for the fine-tuned model (ChromTransfer) while bottom panel displays the sample count for the binary class baseline model (direct training scheme). Note each model and cell line has different ranges of predicted probabilities and therefore different value ranges for each bin.

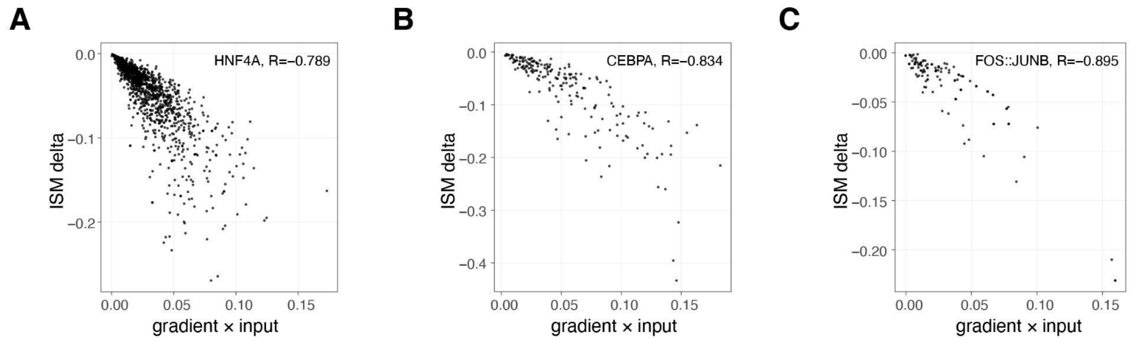


**Supplementary Figure 3: Validation of feature importance scores using in-silico mutagenesis. A-C**: Comparison of gradient × input scores (horizontal axis) and ISM delta scores (vertical axis) associated with predicted TF binding sites of HNF4A (A), CEBPA (B), and FOS-JUNB heterodimer (C). For computational reasons, only true positive predictions were considered. Pearson's correlation coefficients are given in the top right corner of each panel.

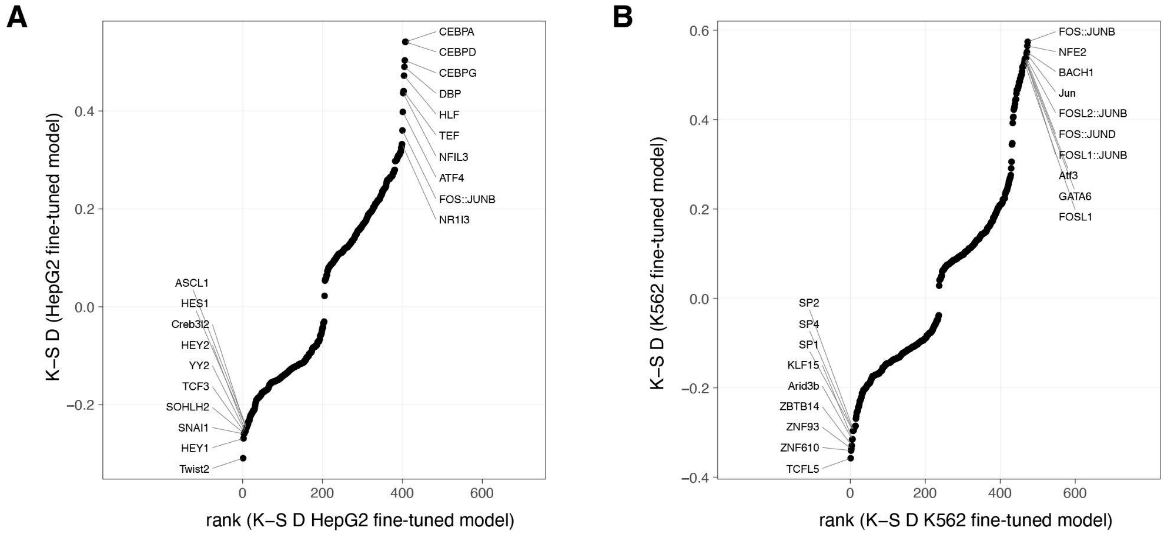


**Supplementary Figure 4: Ranking of feature importance scores reveals TFs important for cell-type specific chromatin accessibility. A-B**: Kolmogorov-Smirnov (K-S) test statistics (D statistics, vertical axes) for feature importance scores (gradient × input) associated with predicted binding site sequences of each considered TF in the HepG2 (A) and K562 (B) fine-tuned models versus their ranks (horizontal axes). Top 10 and bottom 10 ranked TFs are highlighted for each model.

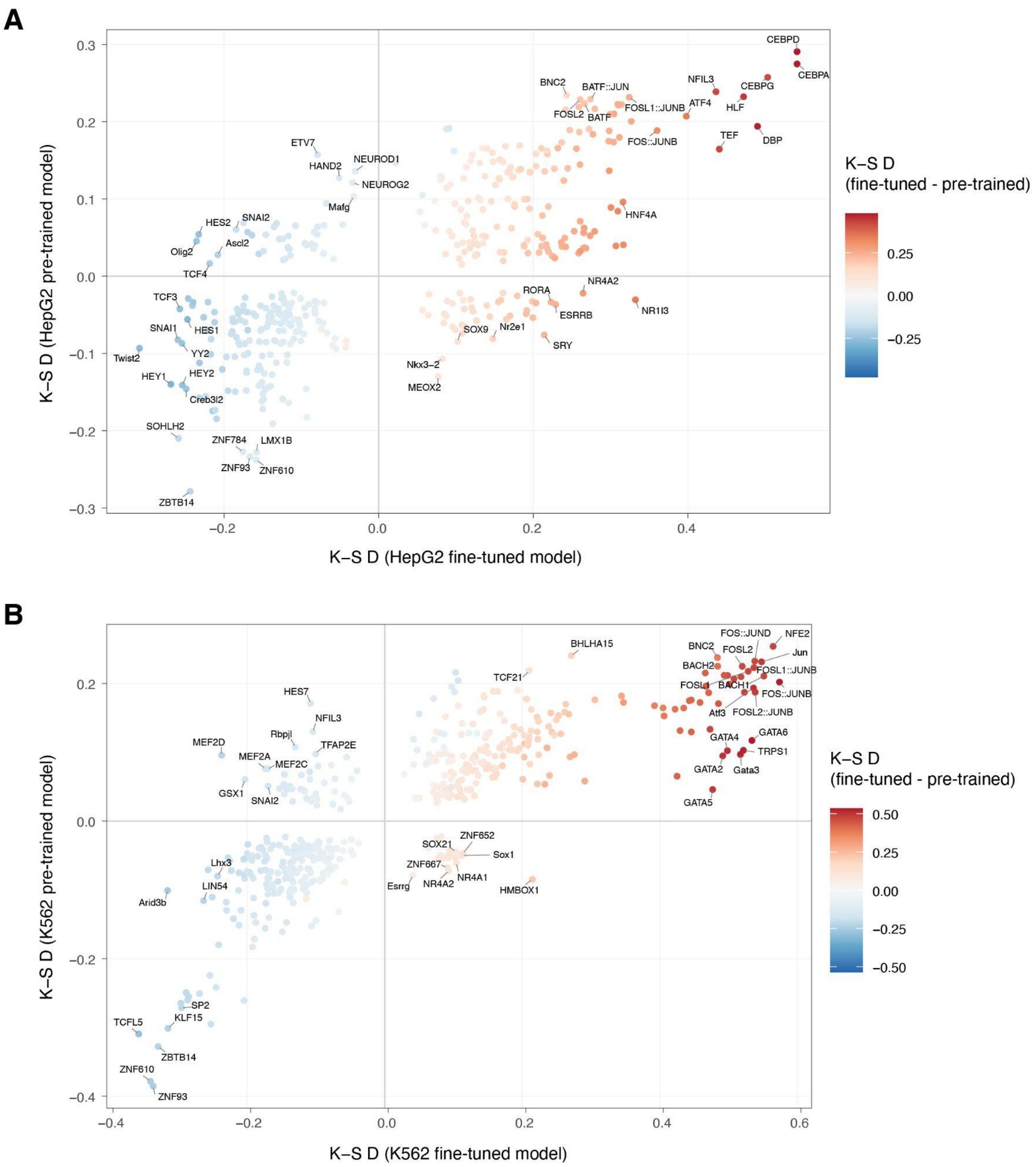


**Supplementary Figure 5: Feature importance analysis reveals how fine-tuning has captured relevant sequence elements for prediction. A-B**: Kolmogorov-Smirnov (K-S) test statistics (D statistics) for feature importance scores (gradient × input) associated with predicted binding site sequences of each considered TF in the HepG2 fine-tuned and pre-trained models (A) and K562 fine-tuned and pre-trained models (B). TFs are colored according to a K-S D statistic calculated from the difference between TF binding site feature importance scores of the fine-tuned and pre-trained models for each cell line. Only TFs with Benjamini-Hochberg adjusted FDR < 0.001 are shown.

### Supplementary Tables

| Cell line | Model | AUROC | F1 overall | F1 neg | F1 pos |
|---|---|---|---|---|---|
| **A549** | **Pre-trained** | 0.74 | 0.32 | 0.31 | 0.33 |
| **A549** | **Baseline (binary)** | 0.72 | 0.8 | 0.91 | 0.26 |
| **A549** | **Fine-tuned** | 0.86 | 0.86 | 0.93 | 0.55 |
| **HCT116** | **Pre-trained** | 0.69 | 0.29 | 0.26 | 0.39 |
| **HCT116** | **Baseline (binary)** | 0.52 | 0.68 | 0.87 | 0.0 |
| **HCT116** | **Fine-tuned** | 0.79 | 0.8 | 0.9 | 0.44 |
| **HepG2** | **Pre-trained** | 0.71 | 0.49 | 0.31 | 0.68 |
| **HepG2** | **Baseline (binary)** | 0.77 | 0.6 | 0.73 | 0.47 |
| **HepG2** | **Fine-tuned** | 0.89 | 0.79 | 0.82 | 0.75 |
| **GM12878** | **Pre-trained** | 0.66 | 0.34 | 0.29 | 0.46 |
| **GM12878** | **Baseline (binary)** | 0.74 | 0.72 | 0.82 | 0.46 |
| **GM12878** | **Fine-tuned** | 0.85 | 0.8 | 0.87 | 0.61 |
| **K562** | **Pre-trained** | 0.66 | 0.24 | 0.22 | 0.33 |
| **K562** | **Baseline (binary)** | 0.82 | 0.81 | 0.91 | 0.39 |
| **K562** | **Fine-tuned** | 0.87 | 0.86 | 0.91 | 0.62 |
| **MCF7** | **Pre-trained** | 0.72 | 0.42 | 0.27 | 0.63 |
| **MCF7** | **Baseline (binary)** | 0.71 | 0.46 | 0.72 | 0.11 |
| **MCF7** | **Fine-tuned** | 0.85 | 0.73 | 0.81 | 0.62 |

**Supplementary Table 1: Predictive performances of the models.** AUROCs, overall and per-class (positive (pos): open chromatin, negative (neg): closed chromatin) F1 scores on the test set for the pre-trained model (prediction of rDHSs) as well as the fine-tuned and binary class baseline models (prediction of cell-type specific chromatin accessibility) of the six considered cell lines.

| | A549 | HCT116 | HepG2 | GM12878 | K562 | MCF7 |
|---|---|---|---|---|---|---|
| **A549** | **0.55** | 0.24 | 0.53 | 0.39 | 0.49 | 0.42 |
| **HCT116** | 0.44 | **0.44** | 0.40 | 0.36 | 0.39 | 0.38 |
| **HepG2** | 0.60 | 0.13 | **0.75** | 0.53 | 0.63 | 0.48 |
| **GM12878** | 0.43 | 0.16 | 0.50 | **0.61** | 0.45 | 0.36 |
| **K562** | 0.51 | 0.16 | 0.51 | 0.40 | **0.62** | 0.34 |
| **MCF7** | 0.53 | 0.23 | 0.61 | 0.51 | 0.51 | **0.62** |

**Supplementary Table 2: Fine-tuning adapts model to cell-type specific predictions.** Positive class F1 scores for each fine-tuned model (rows) using each of the test set data for the six cell lines (columns).

| Cell line | AUROC | AUPRC |
|---|---|---|
| **A549** | 0.61 | 0.24 |
| **GM12878** | 0.75 | 0.56 |
| **HCT116** | 0.62 | 0.38 |
| **HEPG2** | 0.77 | 0.76 |
| **K562** | 0.74 | 0.44 |
| **MCF7** | 0.68 | 0.64 |

**Supplementary Table 3: Performance of the multi-class classification baseline model.** Training and evaluation was performed on the concatenation of fine tuning datasets. Due to different class-balances of the multi-class model, AUPRC values are expected to be lower when compared to the binary classification models.
